# EvoProDom: Evolutionary model of protein families by means of translocations of protein domains

**DOI:** 10.1101/2020.02.23.961532

**Authors:** Gon Carmi, Alessandro Gorohovski, Milana Frenkel-Morgenstern

## Abstract

Here, we developed a novel evolution of protein domains (EvoProDom) model for evolution of proteins, which was based on mix and merge of protein domains. We collected and integrated genomic and proteome data for 109 organisms. These data include protein domain content and orthologous protein families. In EvoProDom, we defined evolutionary events, such as translocations, as reciprocal exchanges of protein domains between orthologous proteins of different organisms. We found that protein domains, which frequently appear in translocation events, were enriched in *trans-splicing* events, *i.e.*, producing novel transcripts fused from two distinct genes. We presented in EvoProDom, a general method to obtain protein domain content and orthologous protein annotation, by predicting these data from protein sequences using the Pfam search tool and KoFamKOALA, respectively. This method can be implemented in other research such as proteomics, protein design and host-virus interactions.

## Introduction

Proteins are built from a set of domains that correspond to conserved regions with distinct functional and structural characteristics (1). In accordance with the domain-oriented view of proteins, domains group together to form domain architectures (DAs), i.e., ordered sequences of domains. Some domains participate in specific DAs, while others are found in diverse DAs. This later property is termed “domain promiscuity” or “domain mobility”. Analyses of domain promiscuity can reveal the mechanisms by which domains are gained or lost (2). Marsh and Teichmann in 2010 (1) described five mechanisms by which proteins gain domains: (i) gene fusion, the fusion of a pair of adjacent genes via alternative splicing in non-coding intergenic regions; (ii) exon extension, the expansion of exon regions into adjacent introns, which encode a new domain; (iii) exon recombination, direct merging of two exons from two different genes; (iv) intron recombination or exon shuffling, insertion of an exon into an intron of a different gene; and (v) retroposition, a sequence located within one gene, which is transposed into a different gene along with a flanking genetic sequence. The mechanism that added a given domain to a protein can be identified from the properties of the gained domain, e.g., the position of the domain in a protein sequence and the number of exons. For example, multi-exon domain gain at the c-termini is due to gene fusion. Additionally, during metazoan evolution, new physical protein-protein interactions (PPIs) emerged consequent to exon shuffling of domains that mediated the interaction (3). An additional work, by Bornberg-Bauer and Mar Albà in 2013 (4), refined and expanded these mechanisms and introduced new concepts such as intrinsically disordered regions, implied links between the emergence of de-novo domains, and de-novo genes (4).

We developed a novel “evolution of protein domains” (EvoProDom) model for the evolution of proteins, which was based on “mix and merge” of protein domains. We collected and integrated genomic and proteome data for 109 organisms. These data included protein domain content and orthologous protein content. In EvoProDom, we defined evolutionary events, such as translocations, as the reciprocal exchanges of protein domains between orthologous proteins of different organisms. We found that protein domains, which frequently appear in translocation events, were enriched in *trans-splicing events, i.e.*, producing transcripts as a slippage of two distinct genes (5). We presented in EvoProDom, a general method to obtain protein domain content and orthologous protein annotation, by predicting these data from protein sequences using the Pfam search tool (6, 7) and KoFamKOALA (8), respectively. This method can be implemented in other research fields such as proteomics (9), protein design (10) and host-virus interactions (11).

## Materials and Methods

The EvoProDom model was based on the data of full genome and annotated proteomes. In addition, the model utilized orthologous protein annotation and protein domain content. Orthologous protein groups were used to group proteins (Refseqs) from different organisms, thereby linking protein domain changes among orthologous proteins with their corresponding groups of organisms. Orthologous proteins were realized as Kyoto Encyclopedia of Genes and Genomes (KEGG) orthologs, namely, KEGG ortholog (KO) (12, 13). Protein domain content was realized as Pfam domains, and this content was associated with proteins. Accordingly, proteins were modeled as a group or list of Pfam domains, and orthologous proteins were modeled as a group of proteins with the same KO number. Both Pfam domains and KO assignments to proteins were predicted from protein sequences alone, using the Pfam search tool (6, 7) and KoFamKOALA (8), respectively. By applying these protein sequence-based methods to obtain protein domain content and orthologous protein annotation, new organisms were easily added to EvoProDom. Moreover, this method is unique to EvoProDom and is applicable to other protein domain-based applications such as proteomics (9), protein design (10) and host-virus interactions (11).

Statistical analysis was performed using R (R: A language and environment for statistical, 3.3.2,2016).

### Data resources

In total, the EvoProDom model was tested on a collection of 109 organisms with full genome and annotated proteomes (Entrez/NCBI (14)). These organisms were grouped according to six categories: (i) 15 fish; (ii) 4 subterranean, 8 fossorial and 21 aboveground animals (15, 16); (iii) 65 organisms with known PPIs (BioGrid version 3.5.173, (17, 18)); (vi) 17 organisms with HiC datasets; (v) 4 cats; and (iv) 15 pathogenic organisms (19). Organisms with HiC datasets were obtained by searching ‘HiC’ in the NCBI GEO database (**Table 1**).

**Table 1.**
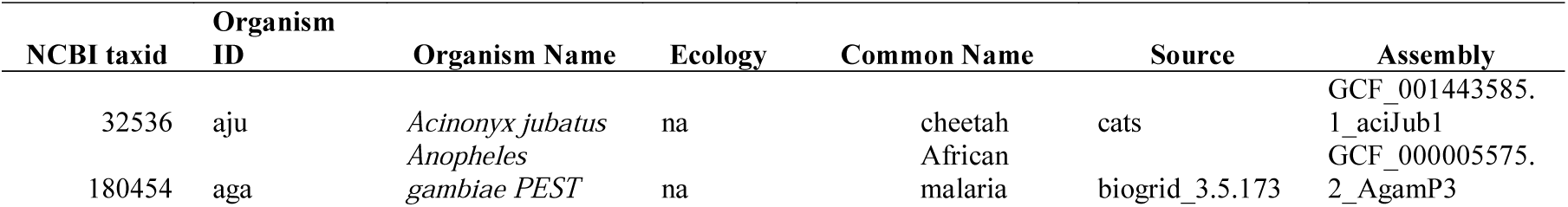
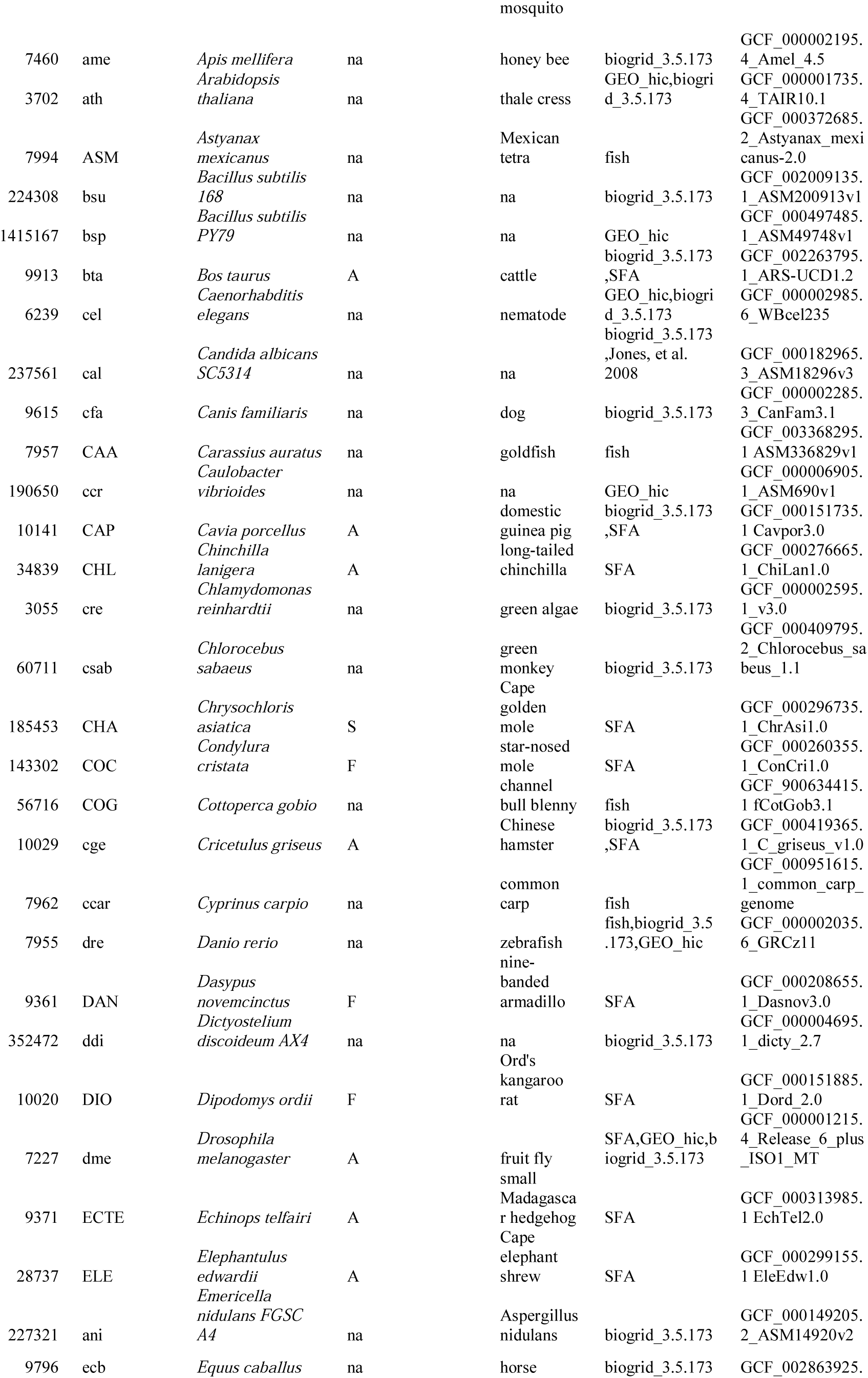
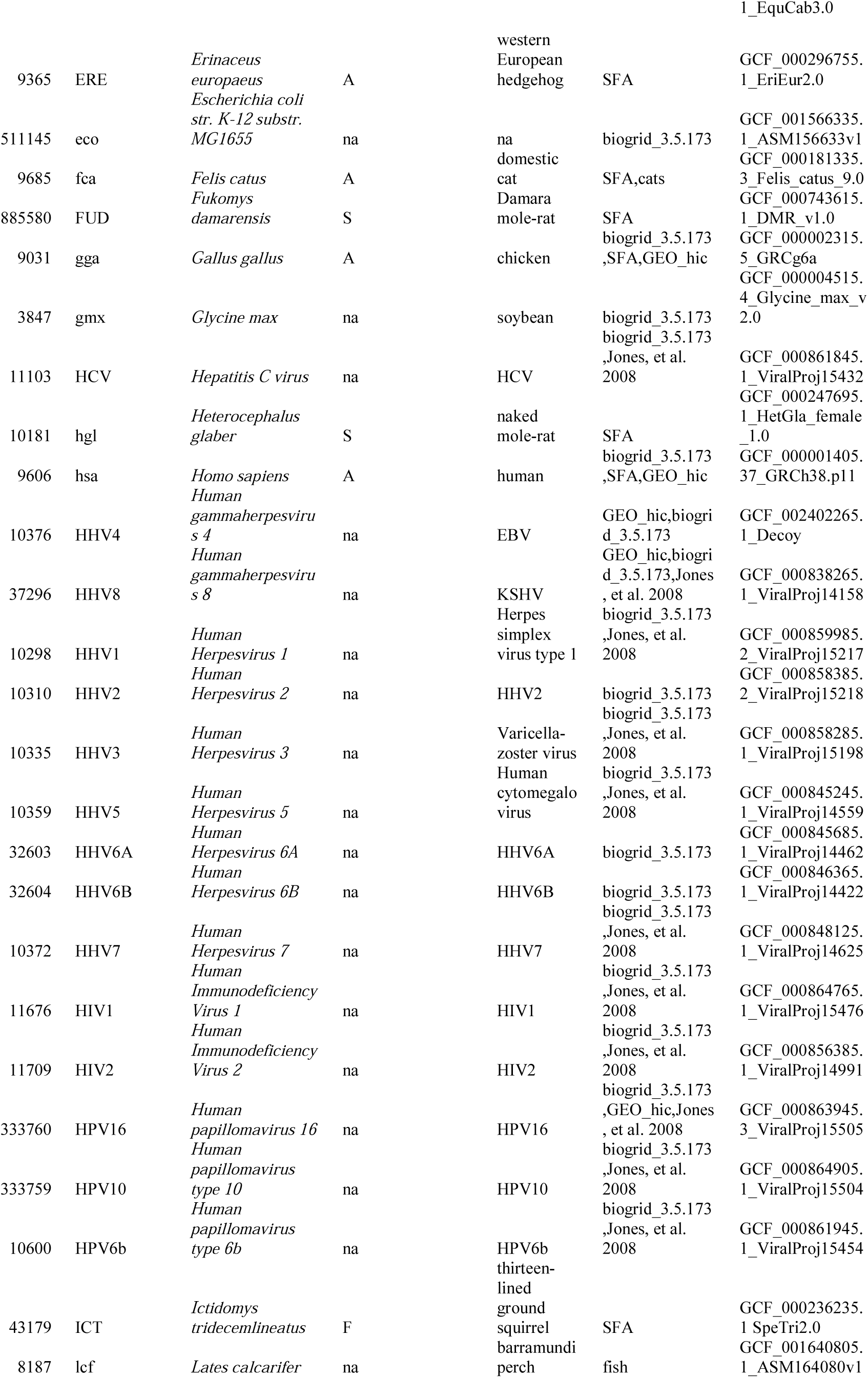
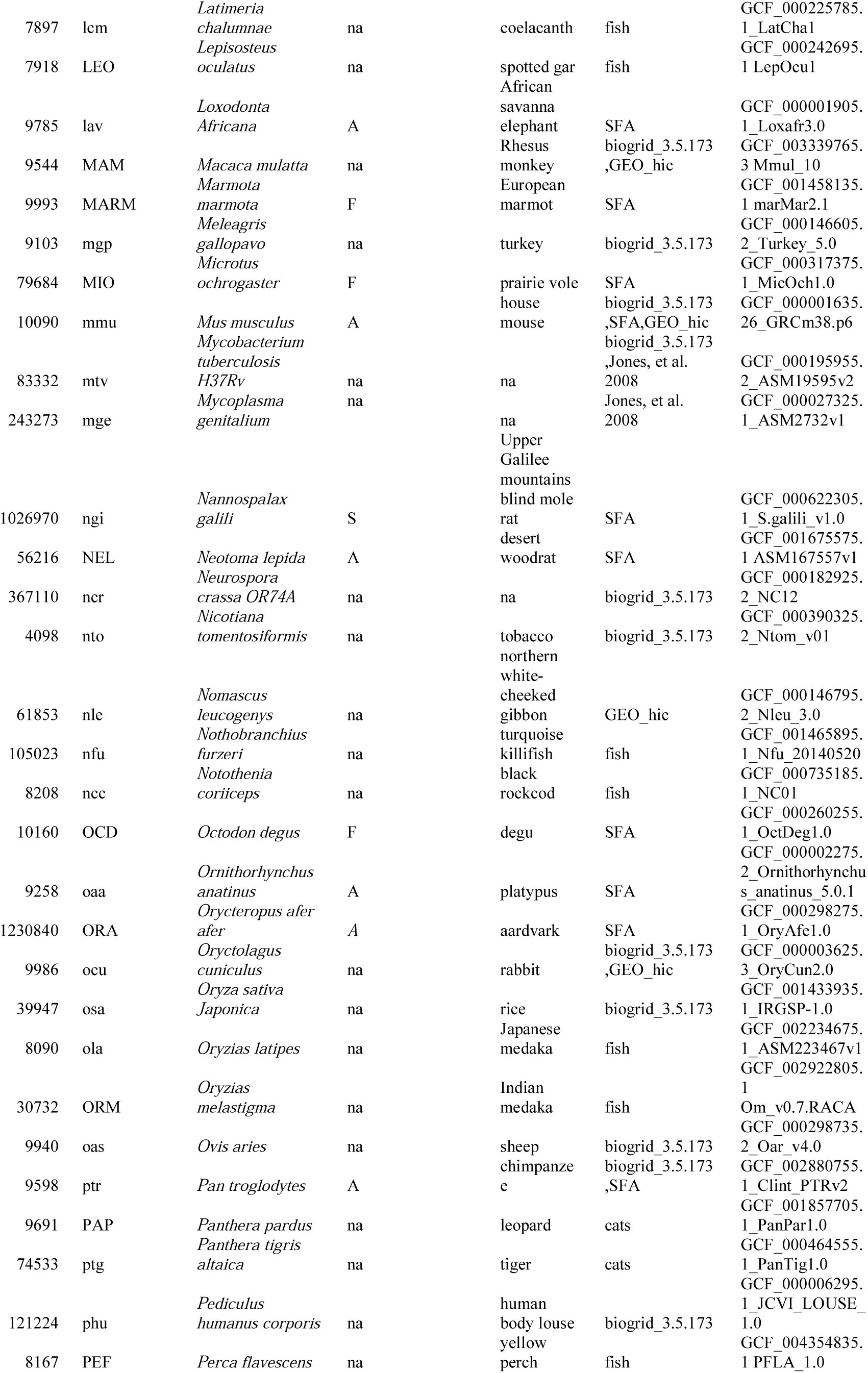
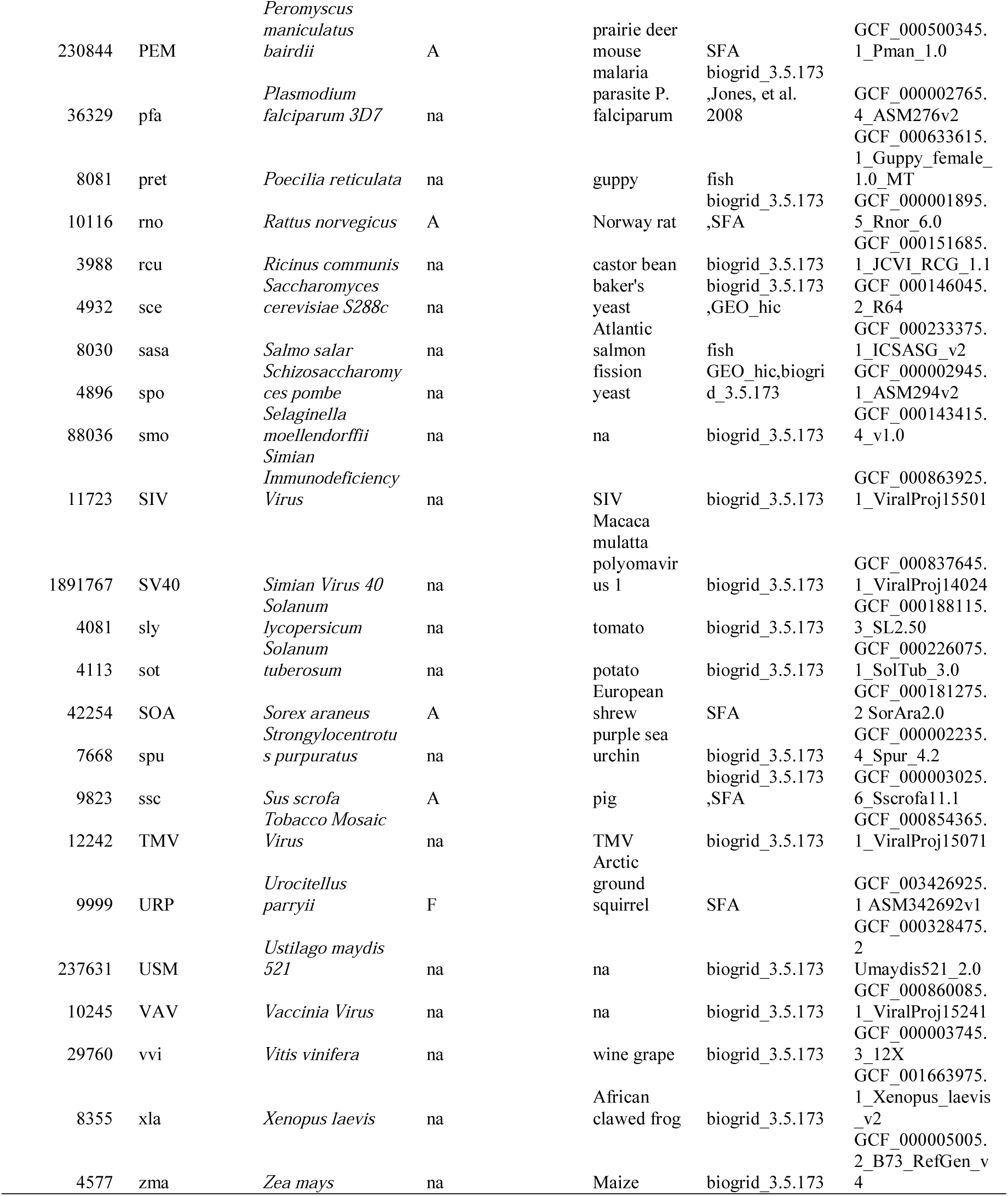
EvoProDom model is applied to an assembly of organisms from diverse taxa. In total, 109 organisms were included in the ensemble and grouped according to six categories: (i) 15 fish; (ii) 4 subterranean (S), 8 fossorial (F) and 21 aboveground (A) animals (SFA) (15, 16); (iii) 65 organisms with known PPIs (BioGrid version 3.5.173, (17, 18)); (vi) 17 organisms with HiC datasets (GEO_hic); (v) 4 cats; and (iv) 15 pathogenic organisms (Jones, et al. 2008) (19). Organisms with HiC datasets were obtained by searching ‘HiC’ in the NCBI GEO database. Taxonomy ID, organism ID, organism name and common name were provided. Additionally, assembly and group classification are indicated. Organism ID is 3-4 letter code were lowercase letter code correspond to KEGG organisms while uppercase letter to organisms not included in KEGG database.

### Orthologous protein annotation

Orthologous annotation was based on Kyoto Encyclopedia of Genes and Genomes (KEGG) orthologs, namely, KEGG ortholog (KO) (12, 13). Proteins were assigned to KO groups, by utilizing KoFamKOALA, a Hidden Markov Model (HMM) profile based search tool (8). Since this tool assigned proteins to KO groups based on protein sequences, in-house script was written to automatically assign proteins to KO groups. In this study, only proteins with KO annotation were used for analysis. Additionally, organism code was generated by selecting 3-4 letters from an organism name in uppercase format (lower case code represents organisms from the KEGG database) (**Table 1**).

### Protein domain detection

Pfam (release 32.0) domains were predicted from protein sequences, using a dedicated HMM-based search tool (6, 7). Therefore, in-house script was written and protein domain content was predicted from protein sequences. Additionally, Pfam domains were members of super families (clan, pfam nomenclature), which we used for our classification. These data were supplemented to the protein domain content.

### EvoProDomDB

Genomic and proteomic data, along with orthologous protein and protein domain content data, were related by shared data. Thus, a relational database, EvoProDomDB, was written in MySQL on MariaDB (10.0.26) for the efficient search engine. The EvoProDom model was implemented and tested on EvoProDomDB. EvoProDomDB was organized with orthologous protein and protein content for the 1,835,600 protein products that were distributed among 23,147 KO groups, containing 17,929 unique Pfam domains. Pfam domains were distributed among 629 super families, and EvoProDomDB integrated data for 109 organisms from diverse taxa. EvoProDomDB was built from six relational tables with common features, e.g., organism id and other features, as shown in the scheme (Figure 1). Relational tables, taxonomy, ko_annotation, clan_domain, pfam_domain provided the annotation data. For example, taxonomy ranks, *e.g.*, species and genus; KO, domain and super family descriptions, respectively.

**Figure 1.**
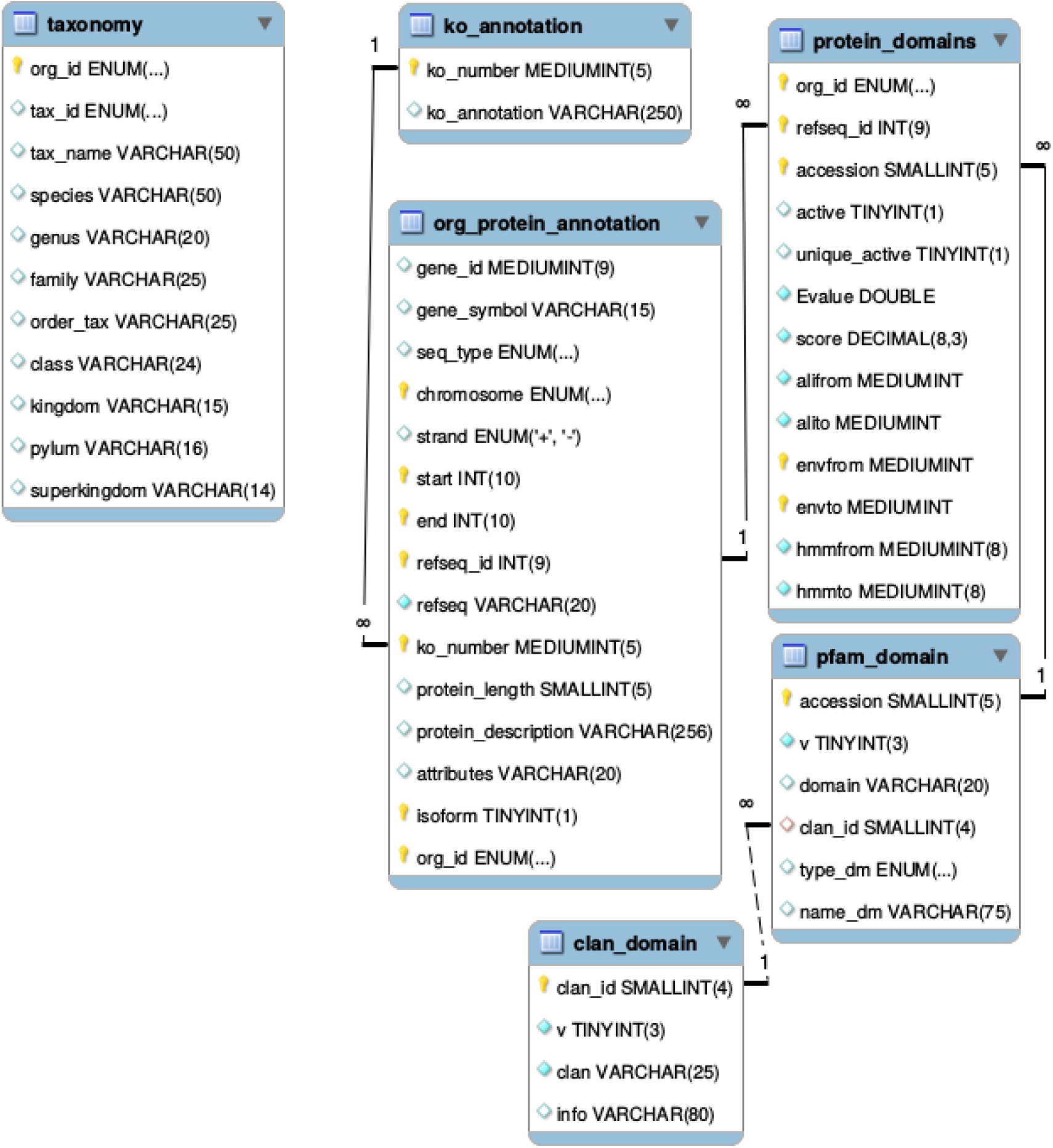
MySQL scheme for EvoProDomDB. Six relation tables were included. Of them, four contained data regarding annotation for taxonomy (taxonomy), KO (ko_annotation,), super families (clan_domain), pfam domain (pfam_domain), such as taxonomy ranks, e.g, species and genus; KO, domain and super family descriptions, respectively. The main relational tables comprise protein, genomic and proteomic data (org_protein_annotation), as well as protein domain content (pfam data) (see the main text for details).

The main relational tables were included for protein genomic and proteomic data (org_protein_annotation), along with protein domain content (Pfam data). Standard genomic and proteomic data were included, for example, gene_symbol, chromosome, strand, refseq_id, a protein length, and a protein description. To these data, the KO number was added (ko_number). Genomic and proteomic data were linked by the longest isoform identification (isoform). Protein domain content included standard Pfam domains as retrieved from Pfam search tool output (6, 7), and calculated data that identified non-overlapping Pfam domains with maximal score (active) delimited by ‘envfrom’ and ‘envto’ coordinates. These corresponded to the largest region where a Pfam domain was predicted to be located within a protein sequence. Among multiple copies of active domains, the highest scoring domain was identified (unique active). To achieve these data, both standard and custom, as well as a construction of EvoProDomDB, in-house bash and perl scripts were written and combined to form a pipeline. The EvoProDom model was implemented as Perl with MySQL quires, to retrieve data from EvoProDomDB and bash scripts. These data sources and databases were summarized in the study workflow (Figure 2).

**Figure 2.**
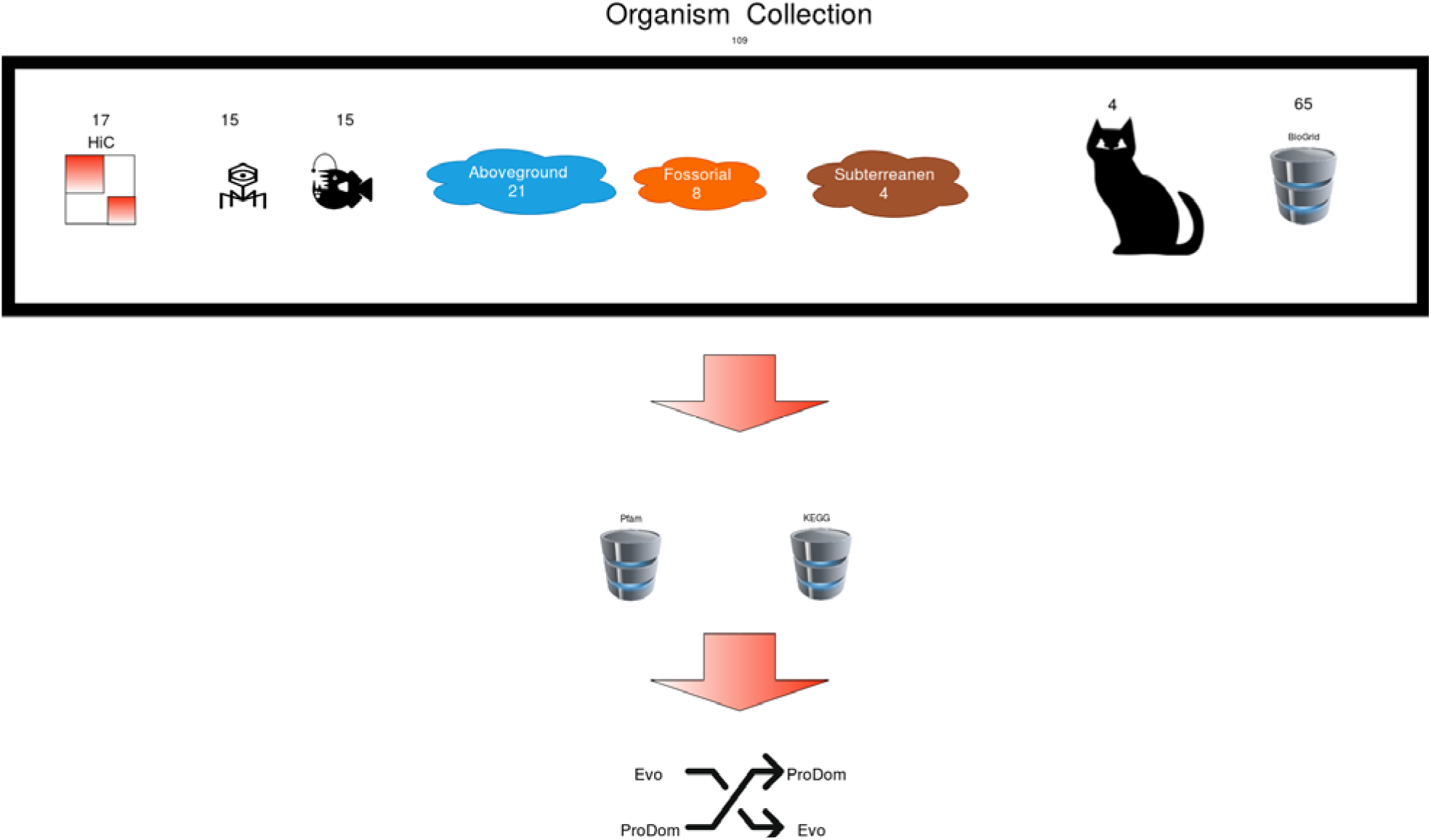
Study workflow: A collection of organisms, 109 in total, was used to implement and test the EvoProDom model. The collection included six categories: (i) 15 fish; (ii) 4 subterranean, 8 fossorial and 21 aboveground animals (15, 16); (iii) 65 organisms with known PPIs (BioGrid version 3.5.173, (17, 18)); (vi) 17 organisms with HiC datasets; (v) 4 cats; and (iv) 15 pathogenic organisms (19). Protein domains were predicted by utilizing Pfam (release 32.0) database along with search tool (6, 7). Orthologous proteins were defined as belonging to Kyoto Encyclopedia of Genes and Genomes (KEGG) (12, 13) ortholog (KO, KEGG ortholog) and assignment to KO group was obtained using KofamKOALA (8).

## Results

### The EvoProDom model

We hypothesized that proteins evolve by means of “mix and merge” or “shuffling” of protein domains, which are distinct functional units (1, 20). The evolutionary model that described protein evolution by means of protein domain dynamics was termed EvoProDom. The EvoProDom model defined and formulated standard evolutionary mechanisms such as translocations, duplications and indel (insertion and deletion) events, which acted on protein domains that were realized as Pfam domains (6, 7). Therefore, proteins, under the EvoProDom model, gained or lost their function based on the presence or absence of respective function-conferring domains. Accordingly, proteins were modeled as sets of protein domains; and evolutionary events, such as translocations, were defined. These describe the gain and loss of particular domains between domain sets or DAs. The KEGG database catalogs diverse taxa and forms groups of orthologous proteins (KO) based on shared function. Thus, all members of the KO group were orthologous proteins (8, 12, 13). Additionally, in the EvoProDom model, proteins were assigned to KO groups (see Materials and Methods). Consequently, translocation events were mapped to groups of organisms according to underlying changes in DA. Thus, evolutionary events, which acted upon domains and manifested as changes in DA were reflected at the organism level. A link between changes at these two levels was therefore established. The EvoProDom model was implemented with and tested on EvoProDomDB (see Materials and Methods). In total, 5,548 translocation events, involving 94 protein super families, excluding an “unknown” super family, were found (*Table 2*). This result indicates the existence of multiple evolutionary translocation events as defined by the model.

**Table 2.** Translocation events per super family (Counts). Super family annotation is provided.

| Super family Id | Super family Name | Counts | Super family Description |
| --- | --- | --- | --- |
| 0010.21 | SH3 | 630 | Src homology-3 domain |
| 0011.26 | Ig | 616 | Immunoglobulin superfamily |
| 0465.3 | Ank | 529 | Ankyrin repeat superfamily |
| 0001.27 | EGF | 414 | EGF superfamily |
| 0361.4 | C2H2-zf | 390 | Classical C2H2 and C2HC zinc fingers |
| 0022.32 | LRR | 282 | Leucine Rich Repeat |
| 0020.25 | TPR | 246 | Tetratrico peptide repeat superfamily |
| 0229.11 | RING | 242 | Ring-finger/U-box superfamily |
| 0186.14 | Beta_propeller | 222 | Beta propeller clan |
| 0221.11 | RRM | 210 | RRM-like clan |
| 9999.0 | Unknown | 210 | null |
| 0159.16 | E-set | 187 | Ig-like fold superfamily (E-set) |
| 0466.3 | PDZ-like | 165 | PDZ domain-like peptide-binding superfamily |
| 0016.22 | PKinase | 164 | Protein kinase superfamily |
| 0266.9 | PH | 141 | PH domain-like superfamily |
| 0023.34 | P-loop_NTPase | 121 | P-loop containing nucleoside triphosphate hydrolase superfamily |
| 0220.12 | EF_hand | 115 | EF-hand like superfamily |
| 0511.3 | Retroviral_zf | 95 | Retrovirus zinc finger-like domains |
| 0271.7 | F-box | 79 | F-box-like domain |
| 0003.21 | SAM | 74 | Sterile Alpha Motif (SAM) domain |
| 0390.4 | zf-FYVE-PHD | 47 | FYVE/PHD zinc finger superfamily |
| 0063.25 | NADP_Rossmann | 37 | FAD/NAD(P)-binding Rossmann fold Superfamily |
| 0357.4 | SMAD-FHA | 37 | SMAD/FHA domain superfamily |
| 0123.18 | HTH | 34 | Helix-turn-helix clan |
| 0680.1 | WW | 34 | WW domain |
| 0167.15 | Zn_Beta_Ribbon | 33 | Zinc beta-ribbon |
| 0006.20 | C1 | 25 | Protein kinase C, C1 domain |
| 0214.13 | UBA | 24 | UBA superfamily |
| 0306.4 | HeH | 24 | LEM/SAP HeH motif |
| 0188.10 | CH | 23 | Calponin homology domain |
| 0459.3 | BRCT-like | 23 | BRCT like |
| 0537.2 | CCCH_zf | 22 | CCCH-zinc finger |
| 0004.20 | Concanavalin | 20 | Concanavalin-like lectin/glucanase superfamily |
| 0072.20 | Ubiquitin | 19 | Ubiquitin superfamily |
| 0033.14 | POZ | 17 | POZ domain superfamily |
| 0154.11 | C2 | 11 | C2 superfamily |
| 0007.18 | KH | 9 | K-Homology (KH) domain Superfamily |
| 0021.18 | OB | 8 | OB fold |
| 0029.20 | Cupin | 8 | Cupin fold |
| 0049.15 | Tudor | 8 | Tudor domain 'Royal family' |
| 0164.13 | CUB | 8 | CUB clan |
| 0172.17 | Thioredoxin | 8 | Thioredoxin-like |
| 0212.9 | SNARE | 8 | SNARE-like superfamily |
| 0392.4 | Chaperone-J | 8 | Chaperone J-domain superfamily |
| 0124.15 | Peptidase_PA | 7 | Peptidase clan PA |
| 0137.15 | HAD | 7 | HAD superfamily |
| 0364.4 | Leu-IlvD | 7 | LeuD/IlvD-like |
| 0575.2 | EFTPs | 7 | Translation proteins of Elongation Factors superfamily |
| 0541.2 | SH2-like | 6 | SH2, phosphotyrosine-recognition domain superfamily |
| 0173.11 | STIR | 5 | STIR superfamily |
| 0192.13 | GPCR_A | 5 | Family A G protein-coupled receptor-like superfamily |
| 0244.9 | PGBD | 5 | PGBD superfamily |
| 0602.2 | Kringle | 5 | Kringle/FnII superfamily |
| 0671.1 | AAA_lid | 5 | AAA+ ATPase lid domain superfamily |
| 0041.13 | Death | 4 | Death Domain Superfamily |
| 0178.16 | PUA | 4 | PUA/ASCH superfamily |
| 0183.14 | PAS_Fold | 4 | PAS domain clan |
| 0642.1 | SOCS_box | 4 | SOCS-box like superfamily |
| 0015.20 | MFS | 3 | Major Facilitator Superfamily |
| 0030.16 | Ion_channel | 3 | Ion channel (VIC) superfamily |
| 0084.13 | ADP-ribosyl | 3 | ADP-ribosylation Superfamily |
| 0198.16 | HHH | 3 | Helix-hairpin-helix superfamily |
| 0497.3 | GST_C | 3 | Glutathione S-transferase, C-terminal domain |
| 0661.1 | Gain | 3 | GPCR autoproteolysis inducing |
| 0027.15 | RdRP | 2 | RNA dependent RNA polymerase |
| 0028.22 | AB_hydrolase | 2 | Alpha/Beta hydrolase fold |
| 0055.13 | Nucleoplasmin | 2 | Nucleoplasmin-like/VP (viral coat and capsid proteins) superfamily |
| 0107.12 | KOW | 2 | KOW domain |
| 0202.11 | GBD | 2 | Galactose-binding domain-like superfamily |
| 0492.3 | S4 | 2 | S4 domain superfamily |
| 0531.2 | AMP-binding_C | 2 | AMP-binding enzyme C-terminal domain superfamily |
| 0005.27 | Kazal | 1 | Kazal like domain |
| 0025.14 | His_Kinase_A | 1 | His Kinase A (phospho-acceptor) domain |
| 0026.20 | CU_oxidase | 1 | Multicopper oxidase-like domain |
| 0070.13 | ACT | 1 | ACT-like domain |
| 0088.16 | Alk_phosphatase | 1 | Alkaline phosphatase-like |
| 0109.12 | CDA | 1 | Cytidine deaminase-like (CDA) superfamily |
| 0110.12 | GT-A | 1 | Glycosyl transferase clan GT-A |
| 0113.13 | GT-B | 1 | Glycosyl transferase clan GT-B |
| 0114.12 | HMG-box | 1 | HMG-box like superfamily |
| 0117.11 | uPAR_Ly6_toxin | 1 | uPAR/Ly6/CD59/snake toxin-receptor superfamily |
| 0125.15 | Peptidase_CA | 1 | Peptidase clan CA |
| 0144.13 | Periplas_BP | 1 | Periplasmic binding protein like |
| 0196.12 | DSRM | 1 | DSRM-like clan |
| 0236.17 | PDDEXK | 1 | PD-(D/E)XK nuclease superfamily |
| 0381.4 | Metallo-HOrase | 1 | Metallo-hydrolase/oxidoreductase superfamily |
| 0426.4 | HRDC-like | 1 | HRDC-like superfamily |
| 0449.3 | G-PATCH | 1 | DExH-box splicing factor binding site |
| 0505.3 | Pentapeptide | 1 | Pentapeptide repeat |
| 0547.2 | GF_recep_C-rich | 1 | Growth factor receptor Cys-rich |
| 0552.2 | Hect | 1 | Hect, E3 ligase catalytic domain |
| 0607.2 | TNF_receptor | 1 | TNF receptor-like superfamily |
| 0630.1 | PSI | 1 | Plexin fold superfamily |
| 0672.1 | p35 | 1 | Baculovirus p35 protein superfamily |
| 0677.1 | GHMP_C | 1 | GHMP C-terminal domain superfamily |

### Mapping of genes to proteins and alternative splicing

EvoProDom combined genomic information (genes) with proteins, and in turn, proteins with Pfam domain composition. In addition, proteins belonging to KEGG ortholog (KO) groups were also included (8, 12, 13). Genes map to more than one mRNA transcript and, in turn, to more than one protein product, i.e., Refseq id. These correspond to isoforms of a gene, which result from alternative splicing, i.e., the inclusion of gene exons. Since, protein domains mostly coincide with exons (1, 3, 5, 20), changes in protein domain content can account for changes in DA as a result of translocation events. Therefore, to avoid confounding effects of alternative splicing, only a longest isoform was used (see Materials and Methods). Therefore, each gene was associated with a single protein product.

### Protein domain content

Proteins are composed of protein domains, which overlap with other protein domains or which have multiple copies. Overlapping domains were undesired as they did not confirm to the linear structure of protein domains within proteins. To address this issue, a single domain (active domain) with a maximal score, as obtained from the Pfam search tool, was determined for each overlapping group of protein domains. This procedure ensured non-overlapping protein domain composition for each protein, consistent with the linear nature of protein domains along a protein sequence. However, multiple copies of such active domains were still present and for translocation events, a unique set of non-overlapping active domains was required. To this end, a similar procedure was applied to multiple copies of active domains, which were referred to as unique active domains. That is, a unique active domain was considered as an active domain with a maximal score among multiple copies of the same domain.

### DA as a basic unit in EvoProDom

According to the EvoProDom model, evolutionary events, e.g., translocations, deletions and insertions, acted on protein domains and involved the DAs, orthologous groups, i.e., KO, and organisms. Hence, EvoProDomDB facilitated in organizing these data according to DA. Briefly, each orthologous group (KO) was partitioned into distinct sets (items), i.e., a list of domains (DA), and matched lists of organisms and proteins. Notably, within these matched lists, duplicated organisms correspond to paralogous proteins. Additionally, for each DA, missing and gained domains were determined from all DAs within a particular KO. From these data, mobile domains and translocation domains, *i.e.*, domains that had undergone all the translocation events, were determined. As a result, we found in total, 5,548 translocation events, involving 94 protein super families, excluding an “unknown” super family (*Table 2*). We identified 2,041 mobile domains of them, 259 had undergone translocation events and 1,782 were involved in indel events (Table *3*).

**Table 3.**
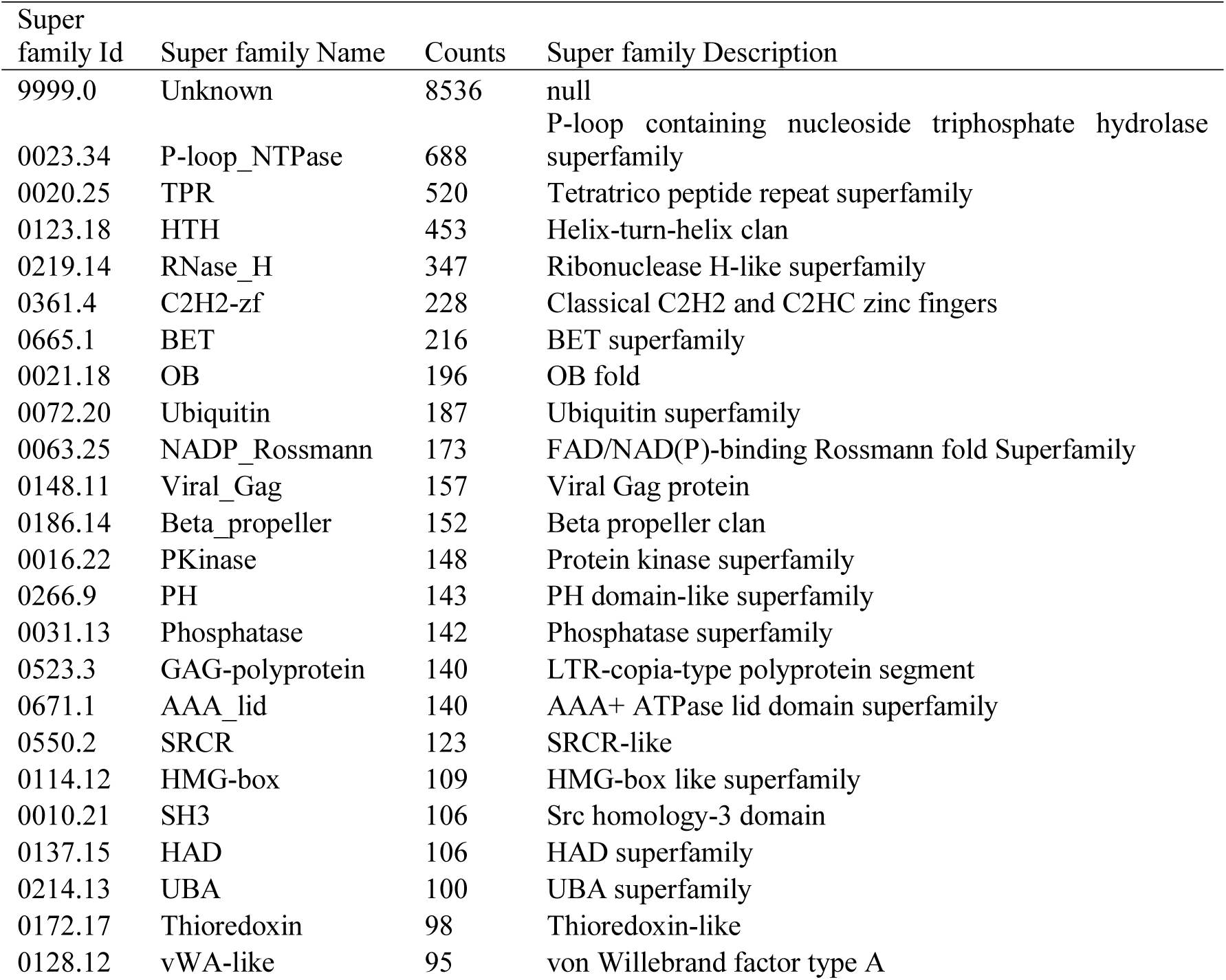
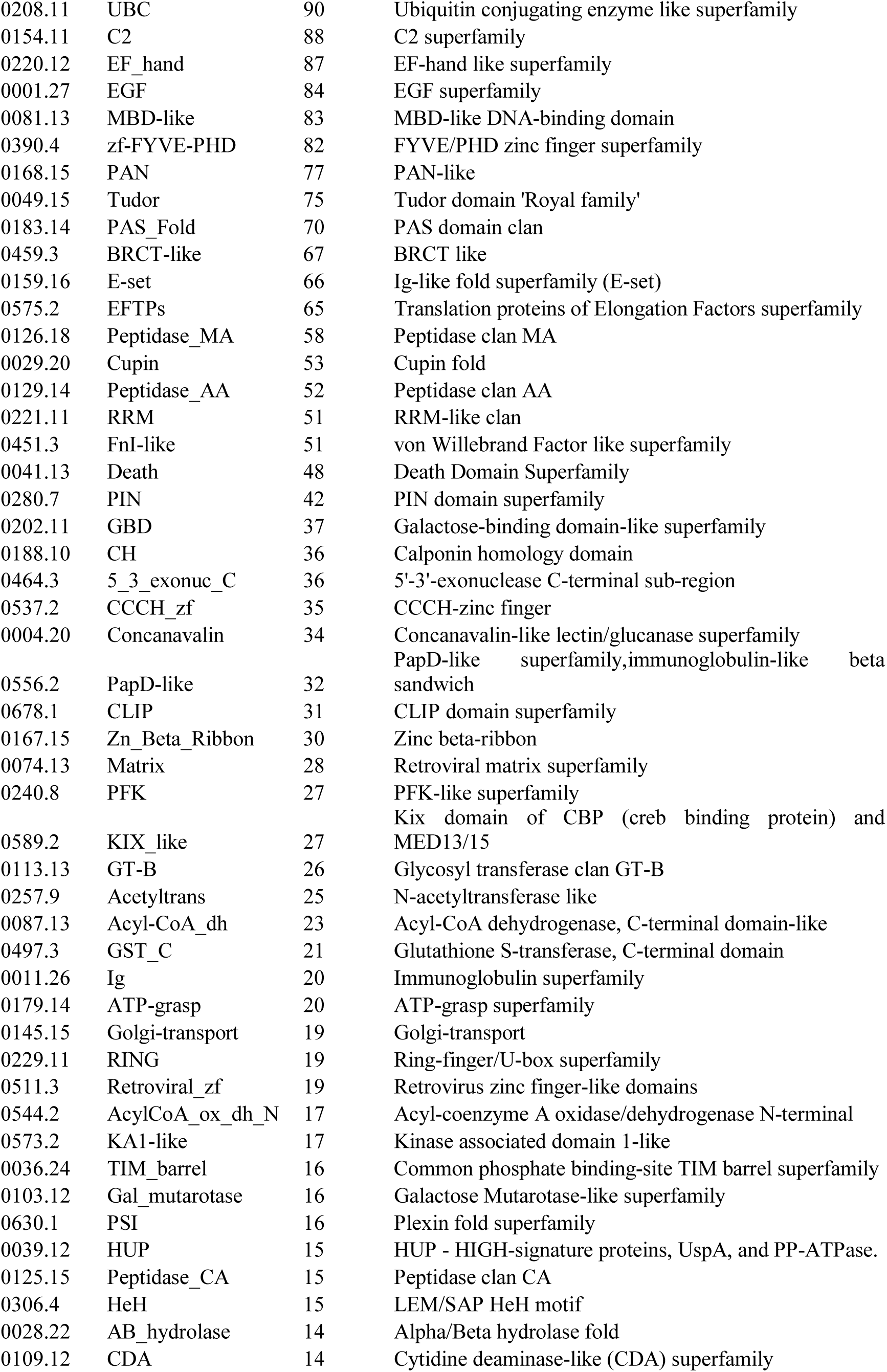
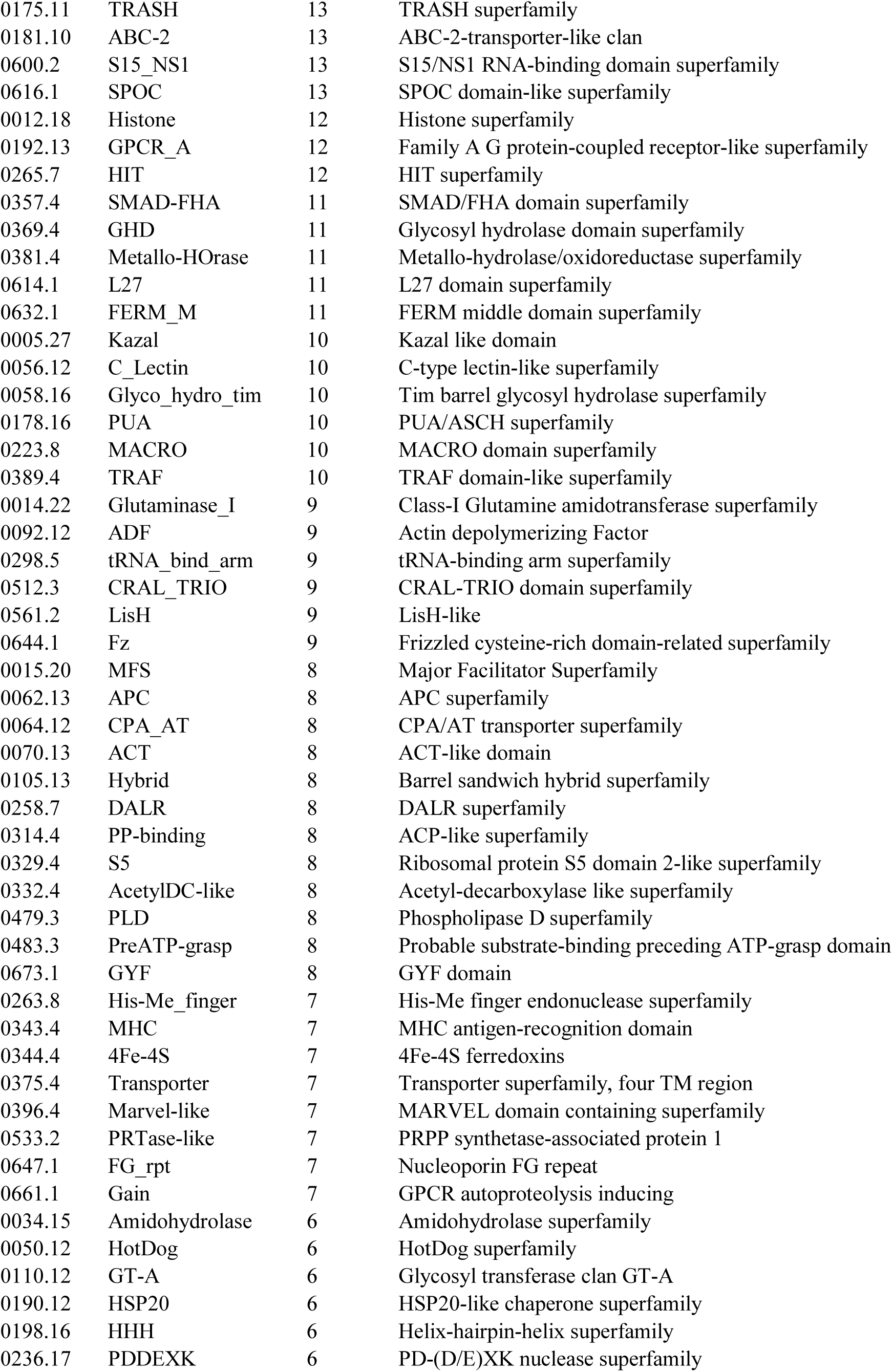
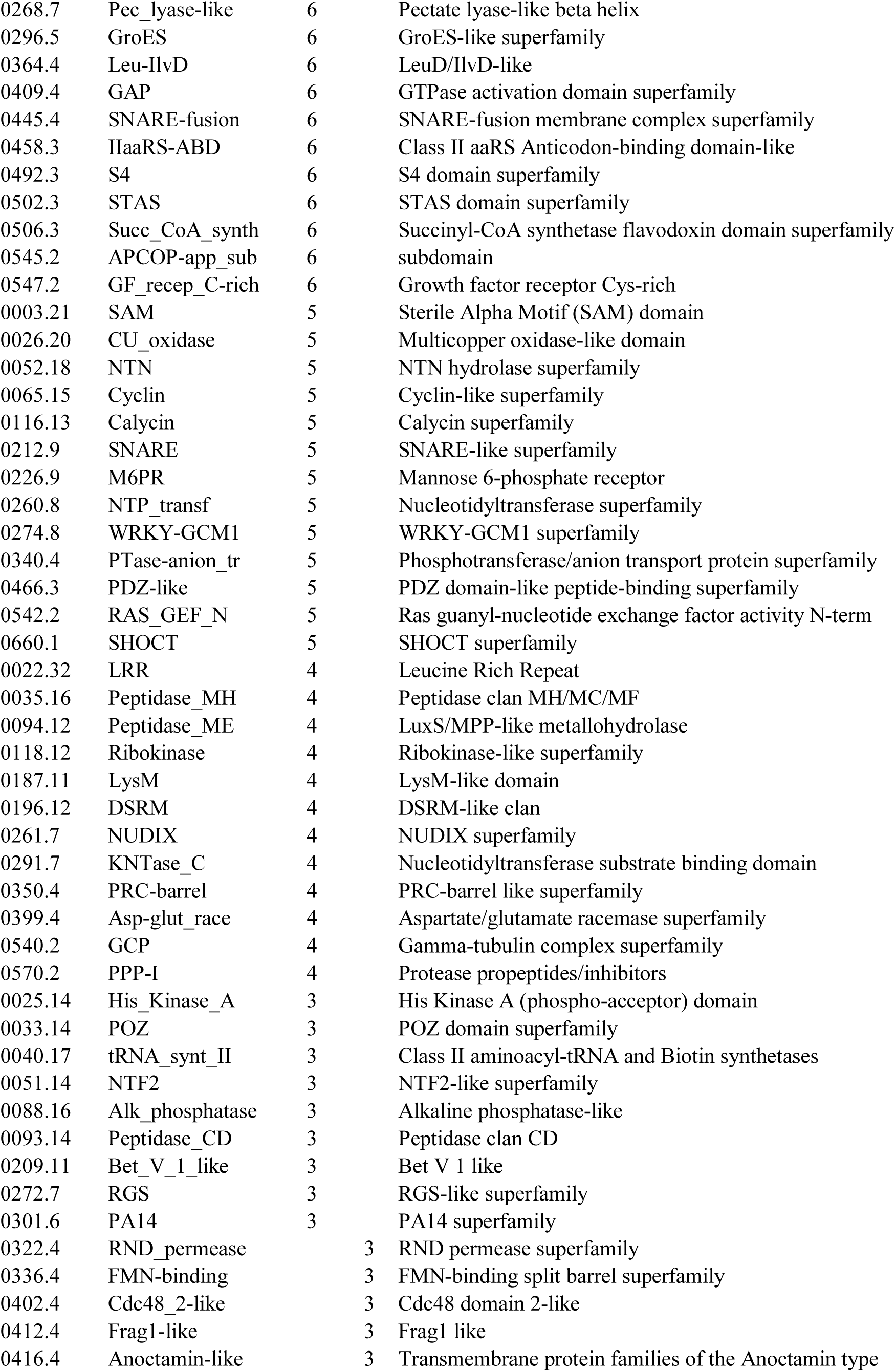
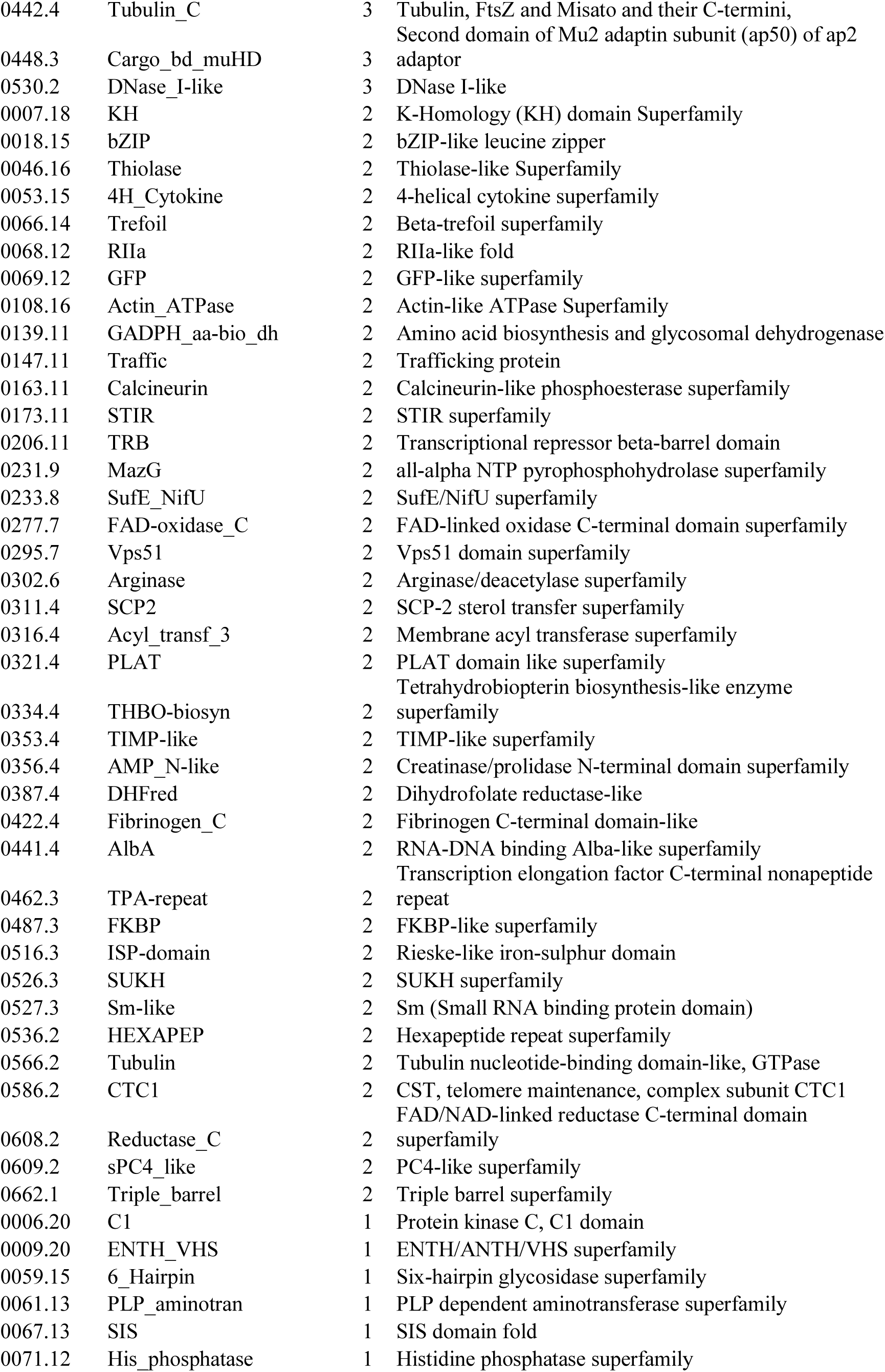
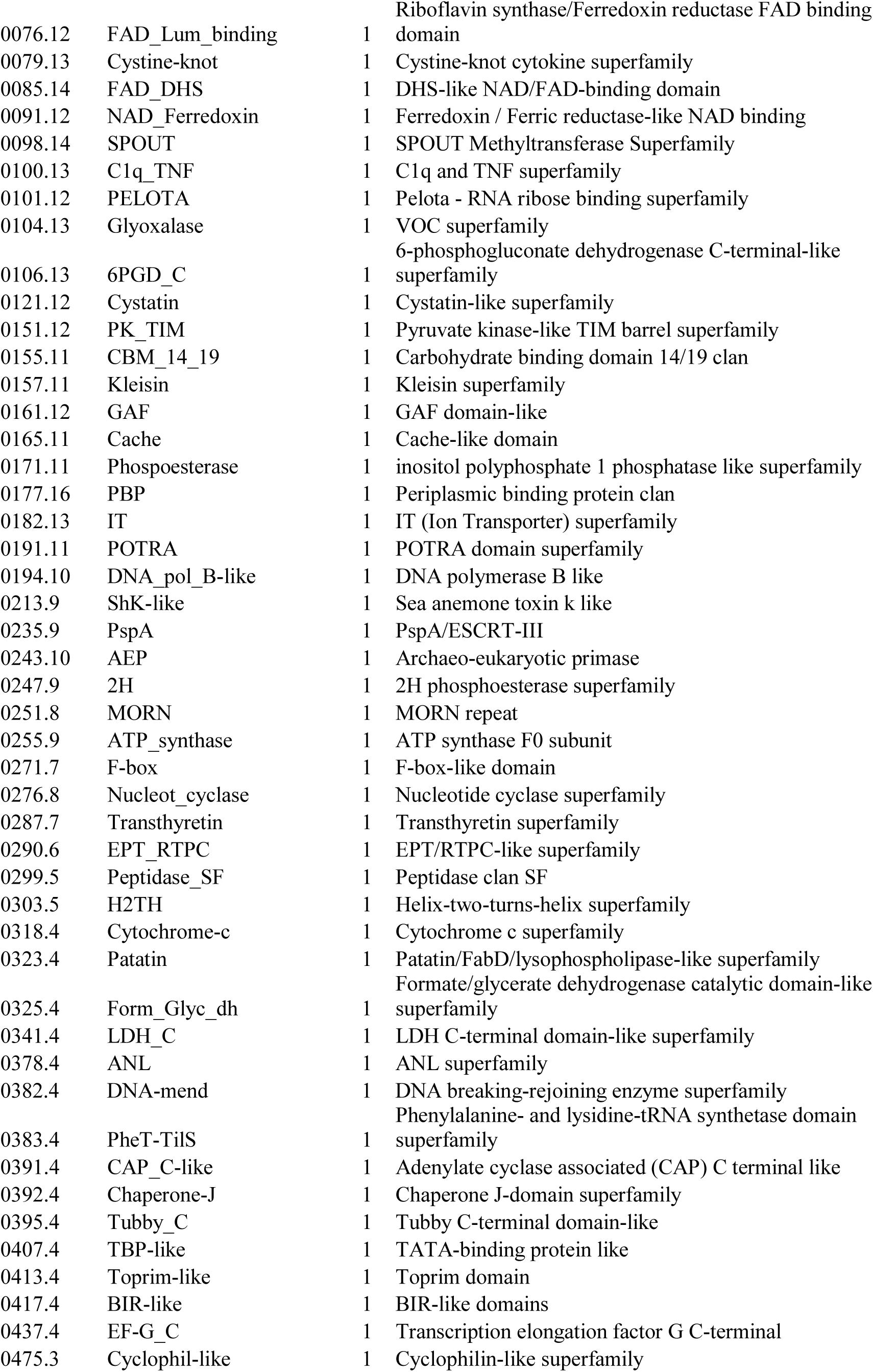
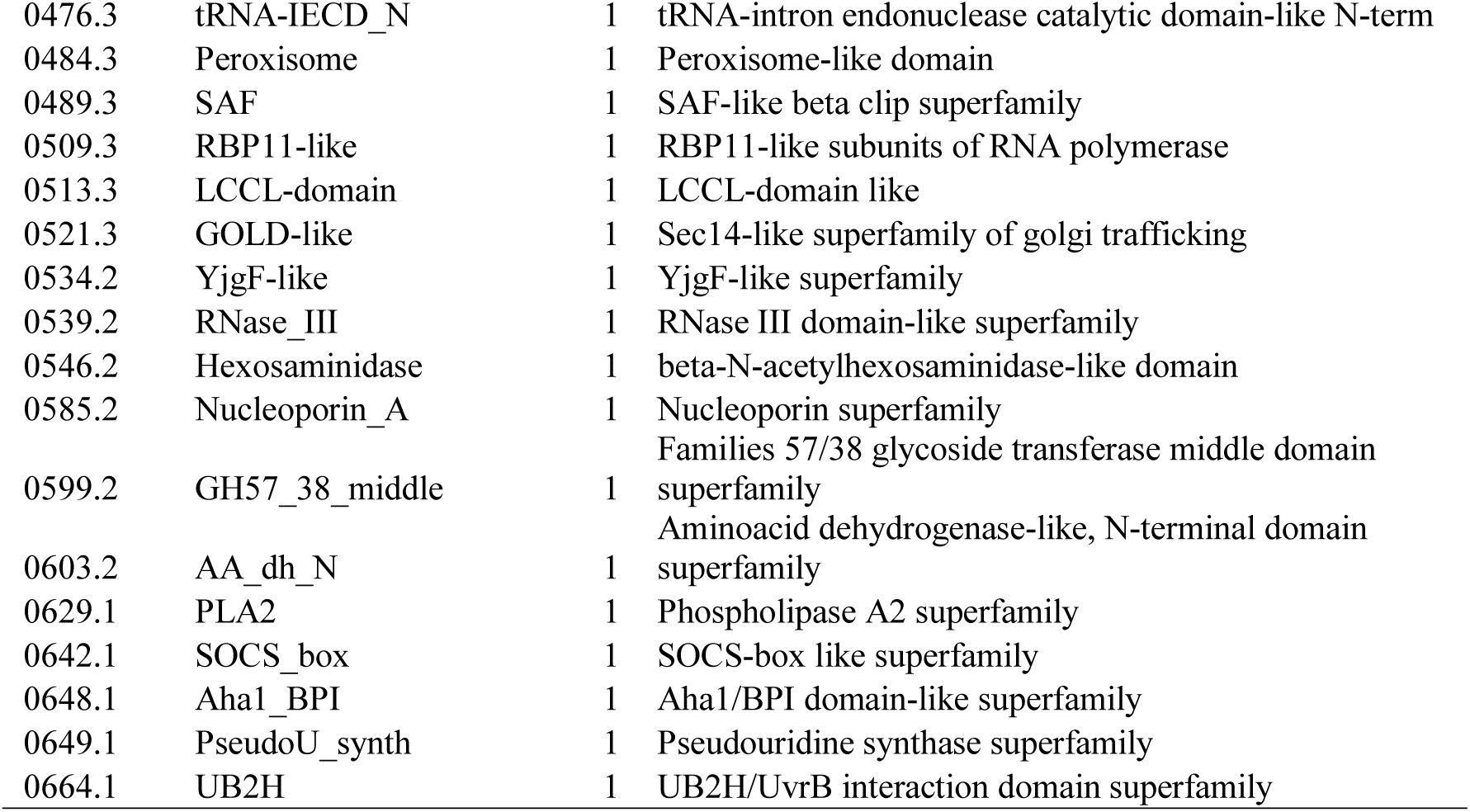
Indel events per super family (Counts). Super family annotation is provided

### Evolutionary mechanism in EvoProDom

#### Implementation of DA

DA, uniquely identified by (ko,item) pair, data were generated from EvoProDomDB, while filtering for active and unique active domains (see Materials and Methods). Each DA included: (i) ko:item; (ii) Pfam domain list; (ii) a list of organisms (org_id); (iii) a list of refseq_ids; (iv) a list of missing domains; (v) and a list of gained domains. Importantly, the list of organisms (ii) and the list of refseq_ids (iii) were matched lists, i.e., the first refseq belonged to the first organism and the second refseq belonged to the second organism, *etc*. Moreover, all other DA information was shared by all organisms and corresponding refseqs, namely, all refseqs were members of the ko group and had similar domain content (item). Gained and lost patterns, (iv) and (v) above, were computed for each KO group among all DAs, i.e., as items. Of note, a minimal number of DA, i.e, items, was two.

Formally, DA, the active domain and unique active domain were defined as follows:

##### Definition: Domain Architecture (DA)

Algorithm: Let *p*_1_, *p*_2_,…, *p*_*n*_ ⊆, were *D* = {*d*_1, *d*_2, …, *d*_*m*} is a set of protein domains and *p*_*i*_ are DA. Grouping of DAs into distinct groups is a partition of *p*_1_, *p*_2_,…, *p*_*n*_

##### Definition: Active domains and unique active domains

Assumptions: protein, *p* {*d*_1_, *d*_2_,… *d*_*m*_}, be DA, *c*(*d*) ∈ ℝ be a score

Algorithm: domain *d* ∈ *p* is an active domain if *c*(*d*) is maximal among overlapping or nested domains.

A unique active domain is the highest scoring active domain among multiple copies of the same domain within p.

#### Translocation and indel events of a mobile domain

Informally, translocations of mobile domains involve gain/loss from/to orthologous proteins from two KO groups, in which the mobile domains were determined by gain/loss patterns within a single KO group. Therefore, a mobile domain was described and defined formally. The main objective of the EvoProDom model was to reflect changes in domain content, namely, at the protein level, with the organism level; and that this would highlight groups of organisms with orthologous proteins, sharing similar patterns of protein domain gain/ loss. Protein domain composition was coupled with organisms by defining mobile and translocation domains. This was based on groups of organisms, and their sizes, with orthologous proteins sharing the same protein domain composition.

Protein domains were contained within orthologous proteins, or the domain missing from a protein, which was based on a number of organisms in each group, *i.e.*, orthologous proteins with and without a particular domain. The mobile domain was defined as follows:

Assumptions: Let *A, B, T* be sets of organisms with proteins in KO group, k, such that *T* = *A* ⋃ *B, A* ⋂ *B =* ø, *O* ∈ *A* {*p* ∈ *O* |*d*_*x*_ ∈ *p*}, *O* ∈ *B* {*p* ∈ *O* |*d*_*x*_ ∉ *p*}. Organisms in *A* contain domain d_x_ whereas organisms in *B* lack domain d_x_.

Algorithm: Unique active domain *d*_*x*_ is mobile between organisms in *A* and in *B* if 4 ≤ |*A*| < |*T*| − 4.

Next, translocations and indel events of mobile domains were defined. Translocations and indel events were mutually exclusive events. Translocation events of mobile domains expanded the patterns of the domain gain and/or domain loss in a single orthologous group to two orthologous groups, such that these patterns generated a reciprocal event. A reciprocal event was characterized by a mobile domain that was gained and lost in the first and second orthologous group, and vice versa (Figure 3). Similar to a mobile domain definition, translocation event criterion was defined for groups of organisms with four or more members. For example, a translocation event of the Pfam domain, PAS_11, is shown in Figure 3. In this translocation event, PAS_11 was present in the KEGG orthologous group number 09095, with correspondence to HIF2A. PAS_11 was absent from the orthologous protein group number 15589 (ARNT2). This gain and loss pattern of PAS_11 was observed among ten orthologous proteins from five different organisms (*A*\*), namely, DIO to ptr, such that each organism had a protein from HIF2A and one from the ARNT2 orthologous group. Reciprocal gain and loss patterns of PAS_11 were observed in orthologous proteins from a second group of organisms (*B*\*), namely, ORO to gaa. A reciprocal gain and loss pattern meant that PAS_11 was absent from the HIF2A orthologous group and gained in the ARNT2 orthologous group. Since these reciprocal patterns of domain gain/loss, between two orthologous proteins, involved two groups of organisms (*A*\* and *B*\*) with more than four organisms each, a translocation event occurred for PAS_11. Note that an additional translocation event occurred for PAS_3 among same organisms and orthologous proteins (Figure 3).

**Figure 3.**
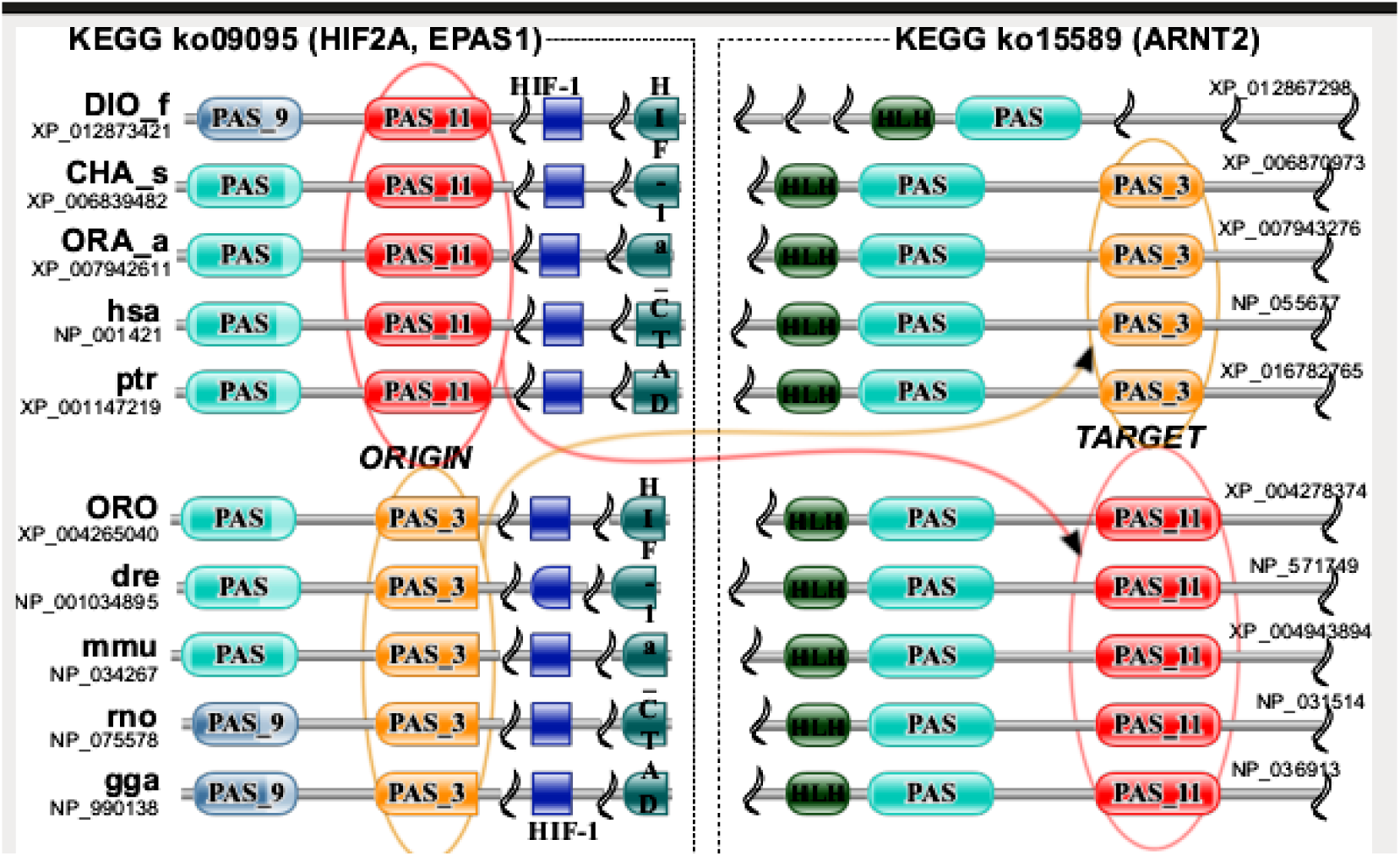
Illustration of translocation event for PAS_11. Red domain (PAS_11) undergone reciprocal a translocation event between two orthologous protein groups K09095 (HIF2A) and K15589 (ARNT2). Accordingly, the red domain (PAS_11) is present in HIF2A and absent from ARNT2 for organisms DIO, etc. while for organisms ORO etc., PAS_11 is present in ARNT2 and missing from HIF2A.

Translocations and indel events were formally defined as follows

Assumptions: Let *d*_*x*_ be a mobile domain between *A*_*i*_ and *B*_*i*_ in *k*_*i*_, where *i* = 1,2, *A*_*i*_, *B*_*i*_ are sets of organisms and *k*_*i*_, are KO groups. Let *A*\* *A*_1_ ⋂ *B*_2_ and *B*\*= *A*_2_ ⋂ *B*_1_.

Algorithm: Mobile domain *d*_*x*_ undergoes translocation if |*A*\*|, |*B*\*| ≥ 4. Otherwise, the indel event has occurred.

#### Duplication of domains

Translocation and indel events were defined for mobile domains; and mobile domains, in turn, were derived from unique active Pfam domains. For duplication events, active domains were used, in order to retain non-overlapping duplicates of Pfam domains (see Materials and Methods). These active domains were calculated for each orthologous protein group, i.e., KO group. Therefore, the duplicate status, “duplicated” or “non-duplicated”, was determined for a particular orthologous KO group and, consequently, varied among orthologous KO groups. Accordingly, the overall duplicate status of a particular Pfam domain was determined by a majority vote from individual KO groups on duplicate status. For example, a Pfam domain’s overall duplicate status was “duplicated”, if the difference between the number of KOs with “duplicated” to the number of KOs with “non-duplicated” status was significant, i.e., 99% percentile of the cumulative sum of the differences. Similarly, overall “non-duplicated” status was determined when considering “non-duplicated” to “duplicated” differences. The duplicate status, namely, “duplicated” or “non-duplicated”, for a particular KO group, was determined based on whether the copy number of particular domains varied or was constant across all DAs respectively. For example, if two copies of a particular domain appeared in one KO while three copies of a domain appeared in a second KO, the domain was determined “duplicated”. However, if a particular domain had the same number of copies within all DA, e.g., two copies, then the domain was considered “non-duplicated”. Duplication was formally defined as follows:

Assumptions: Let *d*_*x*_ be an active domain, *ko* be the KO group with *da*_1_, *da*_2_,…., *da*_*m*_ DAs of active domains. Then *d*_*x*_ is “non-duplicated” in *ko*, if the copy number of *d*_*x*_ is the same in each *da*, otherwise *d*_*x*_ is “duplicated”.

Algorithm: *d*_*x*_ is duplicated if the difference between the number of KO groups where *d*_*x*_ is “duplicated” and the number of KO groups where it is “non-duplicated” is significant (above 99% of the cumulative sum of the differences). Similarly, a non-duplicated domain is defined.

#### Translocation domains were enriched in chimeric transcript

Previously, Frenkel-Morgenstern and Valencia 2012 (5), analyzed enrichment of the domain content for chimeric transcripts combined from two distinct genes. They found that domains were enriched within chimeric transcripts that belonged to the following super families (Super family name): ANK (Ank), EFh (EF_hand), EGF-like (EFG), GTP_EFTU (P-loop_NTPase), IG-like (E-set), LRR (LRR), PH (zf-FYVE-PHD), Pkinase (PKinase), RING (RING), RRM (RRM), SH2 (SH2-like), SH3 (SH3), WD40 (Beta_propeller) and ZnF (C2H2-zf) (5). Of these, EFh (EF_hand), EGF-like (EFG), GTP_EFTU (P-loop_NTPase), IG-like(E-set), Pkinase (PKinase), RRM (RRM), SH2 (SH2-like), SH3 (SH3), WD40 (Beta_propeller) and ZnF (C2H2-zf) were confirmed by RNA-seq data (5). These domains appeared in high copy numbers within proteins, e.g., Ank (21-23) and WD40 (24), as repeats or highly abundant within proteins, e.g., SH3 (25, 26). Therefore, we hypothesized that highly abundant domains might experience a high number of translocation events. To this end, we applied EvoProDom to the assembly of organisms (EvoProDomDB) and identified, in total, 2,041 mobile domains. Of these, 259 had undergone translocation events and 1,782 were involved in indel events (Table *2*, Table *3*). The Pfam domains were classified into super families (6, 7), and translocation events and indel event frequencies were grouped by super families (Table 2, Table 3, respectively). Among the ten most frequent domain super families were SH3 (Src homology-3 domain), Ig (Immunoglobulin superfamily) and Ank (*Table 2*). Likewise, the most frequent super families of mobile domains that were involved in indel events were “Unknown”, P-loop_NTPase and TPR (*Table 3*).

The SH3 super family contains SH3_2 (239 translocations), SH3_1 (198 translocations) and SH3_9 (193 translocations). SH3 (src Homology-3) domains were small protein domains, of approximately 50 amino acids in length (27, 28). They were found in diverse intracellular or membrane-associated proteins (29-31), for example, fodrin and yeast actin binding protein ABP-1. SH3 domains mediate PPIs to facilitate a protein complex assembly (25). The Ig super family contains Ig (219 translocations), I-set (135 translocations), V-set (117 translocations) and Ig_2 (116 translocations). These domains were found in the vertebrate immune system (V-set), cell surface proteins and in intracellular muscle proteins (I-set) (32, 33). Ank repeats super family is comprised of Ank_2 (231 translocations), Ank_4 (184 translocations) and Ank_5 (94 translocations). These domains were found in copies, which form arrays. These repeats were involved in PPIs that regulate cell cycle transition from G1 to S (21-23). This regulation is achieved by the complex formation of INK4 proteins (inhibitors of cyclin-dependent kinase 4) and inhibition of cyclin-dependent kinases 4 and 6 (CDK4/6) proteins (23). These findings show that protein domains enriched in chimeric transcripts undergone many translocations. This supports a connection between chimeric transcripts and EvoProDom translocations. In addition, translocation events for protein domains such as the Pkinase and ubiquitin were found in multiple events and formed a new fusion. Moreover, from each novel transcript, one domain underwent a translocation event (5). Note that both super families with the most and the least number of translocations, SH3 (630) and SH2-like (6), were enriched in chimeric transcripts (Table *2*).

## Discussion

Here, we presented a novel protein evolution model, EvoProDom, which was developed according to the “mix and merge” of protein domains. The EvoProDom model was implemented with and tested on EvoProDomDB. EvoProDomDB consists of genomic and proteome data, along with orthologous protein and protein domain data, for 109 organisms from diverse taxa. In the EvoProDom model, translocations, indel and duplication events were defined to reflect changes in a protein’s domain content among orthologous groups. Moreover, in this model, these changes in protein domain composition were manifested at an organism level. Thus, SH3 was observed as a highly abundant protein domain in translocations, which binds ligands (25, 26) and mediates PPIs (34). Notably, the repetitive domains, such as Ank (21-23) and WD40 (24) appeared in many copies in proteins. These domains fold into 3D structures and generally mediate PPIs (21-24) by means of novel chimeric proteins that form novel PPIs and alter PPI networks of parent proteins (35). Accordingly, these domains, e.g., SH3_2, Ig and Ank_2 and others (see Results), were enriched in multiple fusion event chimeric transcripts (5). As hypothesized, these domains participated in a high number of evolutionary translocation events. The repetition of the domains provides a probable explanation for a high frequency of their translocation events. Fusions were produced as a slippage of two parent genes. These parent genes were truncated at a junction point, resulting in loss of domains. Consequently, the proper function of a chimeric protein is impaired (35). As an example, fusion within a catalytic domain would render the protein non-functional, and such fusion would be selected against. Naturally, due to selection, repetitive domains, which appear in a high copy number, would appear in chimeras at higher frequencies than expected from their shear amount alone, albeit, with lower repeats. Indeed, an average copy number of these domains was reduced in chimeric transcripts (5). In EvoProDom, repetitive domains or abundant domains, e.g., SH3, within KO groups, resulted in a higher number of distinct DAs. This translates into a higher number of (ko,item) pairs (see Materials and Methods). Consequently, repetitive and abundant domains contribute more to the pool of mobile domains from which translocation events were determined, and were thus highly abundant in translocation events. Together, these results indicate that translocation events involving repetitive domains and highly abundant domains rewire PPI networks to achieve adaptive evolution.

The inclusion of new organisms into EvoProDomDB required only full genomes and annotated proteomes. Protein domain content and orthologous protein data were identified from protein sequences using the Pfam search tool (6, 7) and KoFamKOALA(8), respectively. The use of these tools enabled the applicability of EvoProDom to any new organism with a full genome and annotated proteome. Moreover, the usage of these tools in EvoProDom provides a general method to obtain protein domain content and orthologous protein annotation. This method may be applicable to other research fields such as proteomics (9), protein design (10) and host-virus interactions (11). Thus, EvoProDom presents a novel model for protein evolution based on “mix and merge” of protein domains rather than DNA based models. This confers the advantage of chromosomal alterations in evolutionary events.

## List of abbreviations used

DA: Domain Architecture
EvoProDom: Evolution of Protein Domains
PPI: Protein-Protein Interaction
KEGG: Kyoto Encyclopedia of Genes and Genomes
KO: KEGG Ortholog

## Declarations

### Consent for Publication

All co-authors consent to the publication of this manuscript.

## Acknowledgments

M.F.M. is a member of the Dangoor Center for Personalized Medicine and the Data Science Institute (DSI), Bar-Ilan University, Israel.

We would like to thank Dr. Eivatar Nevo for his expertise and helpful comments on the manuscript.

## Author contributions

M.F.M designed the study, supervised it and wrote the paper; G.C, A.G produced the study, verified results and wrote the papers.

